# Purinergic signaling disrupts emergent patterns of multicellular coordination in basal Xenobots

**DOI:** 10.64898/2026.06.04.730190

**Authors:** Thomas F. Varley, Vaibhav P. Pai, Michael Levin, Josh Bongard

## Abstract

The ability of self-organizing systems to display emergent, adaptive capabilities is a fundamental feature of biological life. Understanding the mechanisms by which cells co-ordinate at the micro-scale to produce macro-scale structures and behaviors is a fundamental problem in developmental biology. Moreover, it is an important goal of biomedicine to identify triggers that re-wire the physiological patterns of information flow and control signals. Here, we use a recently developed, synthetic biology platform known as basal Xenobots to explore how patterns of information flow and multi-cellular integration are regulated by chemical signaling pathways. Basal Xenobots are modified, organoid-like systems constructed from embryos, and have previously been shown to display complex patterns of information flows. In this study, we recorded calcium signals before and after exposing basal Xenobots to extra-cellular adenosine triphosphate (eATP), and used statistics from multivariate information theory to explore the differences in global patterns of information flow across the cells in each state. We found that eATP resulted in a dramatic reconfiguration of global information processing dynamics, characterized by a global decrease in multi-cellular co-ordination, reduced information transfer and integration, and a decrease in the global entropy rate of the basal Xenobots. These results provide evidence that purinergic signaling may play a key role in the regulation of multi-cellular self-organization, with implications for a variety of clinical disorders thought to involve aberrant purinergic signaling. These results also suggest the possibility that bioengineers may be able to “tune” the degree of self-organizing capacity in living systems via pharmacological intervention.

## 1 Introduction

A hallmark of biological systems is the capacity for distinct entities (cell, neurons, individuals) to coordinate their activity and generate emergent scales that display their own “collective intelligence” [1, 2]. These emergent systems are capable of sophisticated behaviors, including healing and repair, self-assembly and development, and adaptive responses to dynamic environments. The project of understanding how evolution builds these capacities from basic cellular building blocks has become known as “diverse intelligences” research [3, 4, 5]. A key part of exploring diverse intelligences in biological systems is understanding how groups of cells coordinate their activity and respond to local and global stimuli. This is often framed as “information processing”, as organisms take in signals from their environment and integrate that information into their own on-going dynamics [6]. In this work we use the temporal dynamics of fluorescence signals as a channel to gain insight into information-processing dynamics within collections of cells. Calcium has long been known to play an important regulatory role in coordinating multi-cellular organization across domains of life, such as signal propagation in plants [7], information transfer in oocytes [8], embryonic development [9, 2], healing from wounds [10] and sunburn [11], and neural signaling [12].

Understanding self-organizing processes in living systems is complicated by the fact that emergent behaviors are typically assumed to be the result of active natural selection over evolutionary timescales [13]. This leads complex, multi-scale dependencies where selection for macro-scale structures constrains the behavior of micro-scale structures, while simultaneously, the capabilities of the micro-scale cells inform and constrain the kinds of macro-scale structures that are possible. This mixture of downward and upward-causation creates reciprocal interactions between scales, making it difficult to untangle “bottom-up” and “top-down” emergent structure. Here we turn to synthetic biology - specifically “basal Xenobots”, which are artificial constructs with no macro-scale evolutionary history, and so any observed patterns must be the result of genuinely bottom-up self-organizing processes playing out at the cellular level. The goal is to identify the effects of a perturbation at the micro-scale (in this case, exposure to extracellular ATP) on the innate self-organizing capacity of the cells in a context that lacks the downward-causal pressures built in by evolution.

The Xenobot platform (autonomously moving biobots derived from Xenopus cells) provides an excellent model system that is a minimal, synthetic, nonneural, and multicellular system, to take the first steps towards applying quantitative information-theoretic measures to understand and quantify complexity. The Xenobot platform constitutes a multitude of autonomously motile, self-assembling constructs that can be formed from single or combination of tissues, actuated with either ciliary motion or muscle activity [14, 15, 16, 17, 18]. They can be constructed into multitude of shapes, show self-healing, kinematic replication, transcriptional plasticity, and capability to hold memories of past experiences [13]. Here we use basal Xenobots (spherical, single tissue, no synthetic circuits, no scaffolds, no sculpting, no engineering) to test the information-theoretic measures in the simplest possible Xenobots and their responses to external stimuli.

A fundamental challenge to this work, however, is the problem of quantitatively describing the complex, multi-scale relationships between “parts” and “wholes” in self-organizing systems. Biological systems are high-dimensional, non-linear, active matter [19]; cells are both part of a higher-order collective, and semi-autonomous agentic systems of their own. This scale-free structure challenges standard analytical approaches based on linear models and assumptions of reductionism and independence that have historically dominated life sciences. In response, information theory (a branch of mathematics naturally suited to learning dependencies from complex systems), has emerged as a powerful formal framework for understanding complex systems [20]. Information theory provides a natural scaffold for studying complex systems, as it can easily represent both dynamic dependencies and multi-scale part/whole relationships using a well-developed formal language that describes how knowledge of one “part” changes our uncertainty about the state of another “part”.

Recently we showed that basal Xenobots, despite their non-neural structure, display sophisticated patterns of multicellular information processing, including emergent, meso-scale structure, information transfer, and information integration [21, 22]. We hypothesized that these patterns represented a functional scaffold that reflected the self-organizing capacity of the individual cells into a coherent system. That work, however, could not directly test whether the observed pat-terns of information reflected specific biological processes related to self-organization, or if they were merely epiphenomena. Prior work on perturbing Xenobots showed that physically damaging their structure via stabbing resulted in altered cell-cell functional connectivity networks [21], but this is an extreme intervention stabbing physically severs connections between cells and so changes in functional connectivity are somewhat obvious and do not necessarily reflect a change in the biological processes of self-organization and active matter. To test whether information dynamics provided insight into specific biological processes relevant to self-organization, we took inspiration from computational neuroscience, where psychoactive drugs and neuroimaging analyses have long been used to probe the link between biological structure and dynamic function. We re-analyzed recently reported data on the effects of extracellular ATP (eATP) exposure on calcium signaling dynamics in basal Xenobots [13]. Xenobots were imaged before exposure to eATP, and then again four hours post-exposure. Drawing from prior neuroscience studies [23, 24, 25, 26], we then analyzed the patterns of cell-to-cell signaling using a battery of statistical measures of information-processing dynamics, aiming to create a comprehensive portrait of how purinergic signaling alters multicellular coordination.

The information dynamics framework uses tools from information theory to decompose “computation” in complex systems into distinct components that reflect increasingly higher-order structures and very roughly map onto aspects of digital computation [6, 27]. The first, and lowest-order information dynamic is the active information storage [28, 29], which quantifies the amount of information that a cell’s past state discloses about its own future. It can be seen as a measure of memory, analogous to the autocorrelation function. The second measure is the pairwise information transfer (typically formalized by the transfer entropy [30, 27], formalized as the information that the past of one cell discloses about the future of another cell. For Gaussian systems, the transfer entropy is directly related to the Granger causality [31], a popular measure of time-directed dependency in biology and economics. Intuitively, it can be understood as a measure of time-directed information “flow”, in contrast to the active information storage, which is a measure of time-directed “memory”. The final type, information modification, quantifies the amount of novel information generated when multiple incoming “streams” of information are processed by a target. Here, we operationalize it using the integrated information [32, 33], which measures the time-directed information present in the joint-state of a set of variables that is not learnable from any subset (this is often called “synergistic information”).

Collectively, these results paint a portrait of how purinergic signaling alters different aspects of information processing in the basal Xenobots, at multiple scales. Changes in these information flows reflect alterations to the self-organizing capacity of the biobots in response to a known, biochemical perturbation, and point the way towards a rigorous theory of structure/function coupling and emergence in living systems.

## 2 Materials and methods

### 2.1 Animal Husbandry

For all experiments, Xenopus Laevis embryos were fertilized in vitro as per established protocols [34] and reared in 0.1X Marc’s Modified Ringer’s (MMR) solution. Embryos were maintained at 14°C, staged according to Nieuwkoop and Faber [35], and microinjected at 2-cell stage with ectodermal explants excised at stage 9.

### 2.2 Microinjection

Microinjection was performed using standard protocols [34]. mMessage mMachine kit (Thermo Fisher Scientific, AM1340) was used to synthesize capped synthetic mRNA from a linearized DNA template and stored in nuclease-free water at -80°C. Embryos were put in 3% Ficoll solution and synthetic mRNA was microinjected with a pulled glass capillary needle. Both blastomeres at 2-cell stage were injected for ubiquitous expression. 1-2 ng of mRNA was microinjected into each blastomere at the center of the animal pole. Embryos were left to heal for 30 mins in Ficoll followed by replacing half of Ficoll with 0.1X MMR with 15-minute incubation at room temperature. Embryos were then washed five times in 0.1X MMR to remove all Ficoll solution. Unhealthy or damaged embryos were discarded. Embryos were incubated at 14°C. GCaMP6s (a calcium dynamics reporter) mRNA was microinjected.

### 2.3 Basal Xenobot preparation

Stage-9 Xenopus embryos were used for generating basal Xenobots (no sculpting, no engineering, no scaffolding, and no mixing of tissues) as previously described [14]. Briefly, stage-9 Xenopus embryos were moved into a 1% agarose (made with 0.75X MMR) coated petri dish containing 0.75X MMR. Embryos were devitellinized and ectodermal explants (animal pole progenitor cells – animal cap) were cut according to established protocols [36, 37, 38, 39]. Over 2 hours explants were allowed to close in on themselves and form round spherical structures. These were then washed five times with 0.75X MMR and moved to a new 1% agarose coated petri dish and incubated at 14°C for seven days with daily cleaning. Over seven days, the rounded explants transform into autonomously motile atypical/synthetic epidermal entities - mature basal Xenobots, used for all experiments [14, 40, 41, 42, 43]. Basal Xenobots showing uniform and robust fluorescent signal were used for calcium-dynamics imaging.

### 2.4 ATP treatment

Adenosine triphosphate (ATP) (Sigma-Aldrich, A1852) was dissolved in 0.75X MMR at a stock concentration of 100 mM and aliquoted into single use portions and stored at -20°C. Each aliquot was thawed immediately before the experiment and 10 *µ*L of stock solution was added per 4 mL of 0.75X MMR to achieve a final concentration of 250 *µ*M.

### 2.5 Calcium imaging and preprocessing

Bespoke chambers [13] were used to image GCaMP6s-expressing Xenobots using Leica Stellaris 8 confocal microscope with 25X objective. Each Xenobot was imaged for 15 mins with a capture rate of one Z-stack projection every 10 seconds. GCaMP6S and LifeAct-mCherry were imaged in parallel with separate excitation wavelengths and separate detectors, although only GCaMP6S is analyzed herein. Preprocessing videos involved extracting just the green channel, and registering the frames using TV-L1 optical flow alignment [44] implemented in the OpenCV computer vision package [45]. Finally, after registration, the individual cells were segmented using the Cellpose [46] package, following [21]. From those segmented cells, the average calcium signal for all pixels within the cell were averaged to create a cell-level calcium signal. Finally, all cell-level calcium time series underwent z-scoring and global signal regression to remove artifacts attributable to global changes in luminance or visual field. Basal Xenobots were imaged both before any exposure to ATP stimulus, and 4 hours after ATP exposure. See Figure 1.

**Figure 1:**
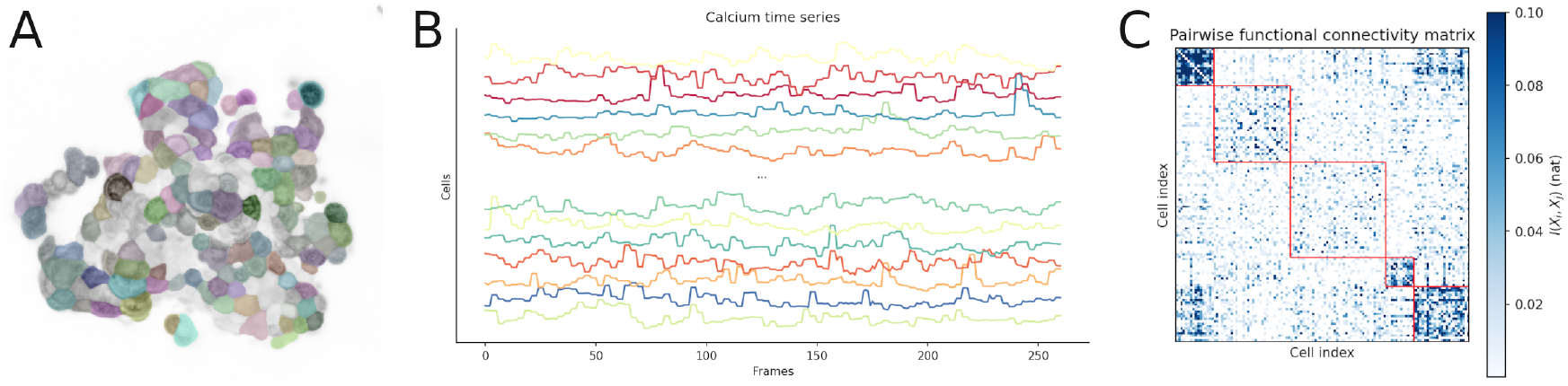
Statistical network inference pipeline. **A:** A video of a basal Xenobot is segmented at the cell level using the Cellpose neural network system [46]. **B:** The calcium intensity signals from all pixels within a cell are averaged to construct a time series for each cell. **C:** From this multivariate time series, functional and effective connectivity networks can be constructed: here is a pairwise, mutual information functional connectivity network as an example. The network was clustered using a multi-resolution consensus clustering algorithm [49] to highlight the non-trivial structure in the information patterns.

### 2.6 Null model

The calcium data analyzed here is known to be highly autocorrelated [22], which presents a confound, as autocorrelation can artificially inflate the apparent magnitude of cellular interactions [47]. To address this, we correct all estimates by subtracting off (or otherwise normalizing) the raw estimates with the expected value of a null distribution computed from circular-shifted surrogate data [48]. Briefly, for each measure of interaction, we first compute it from the true data, and then we re-compute it a large number of times from data that has had each channel shifted randomly forward or backwards in time. Shifts for each channel were chosen randomly from a uniform distribution on the range [*− T/*2, *T/*2], where *T* is the number of frames in the data. The circular shift null preserves all first-order features of the data (autocorrelation, spectral properties, etc), while disrupting any interactions. By normalizing the empirical values by the expected null values, we can remove any apparent dependency that is actually attributable to first-order features of the data, leaving only the “true” interactions.

### 2.7 Information-theoretic analyses

Information theory has become something of a *lingua franca* for the study of complex systems, as it provides a natural formal framework with which to characterize multipartite interactions between elements [20]. Here, we revisit many of the analyses described in our previous paper on basal Xenobots [22] to establish that these methods are sensitive to biologically-relevant differences in biological structure and function.

The basic object of study in information theory is the Shannon entropy. For a potentially multidimensional random variable **X** which selects states from a support set **𝒳** according to probability distribution *P* (**x**), the entropy is:

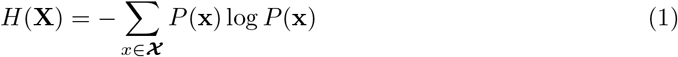

For real-valued random variables, the entropy can be estimated using parametric Gaussian assumptions. The entropy of a *k*-dimensional Gaussian with a covariance matrix ***S*** is given by:

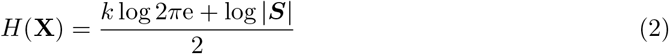

The pairwise dependency between cells can be estimated with the Shannon mutual information:

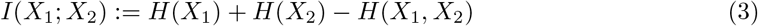

For Gaussian random variables, mutual information between two one-dimensional channels is closely related to the Pearson correlation coefficient *r*:

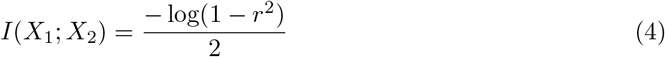

While for two multivariate random variables **X** and **Y** with covariance matrices **S**_**X**_ and **S**_**Y**_, it is given by:

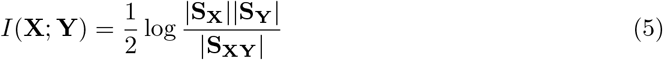

#### 2.7.1 Cellular active information storage

The simplest measure was the active information stored in each cell’s calcium signaling time series [6].

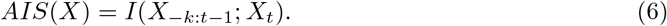

Here the parameter *k* refers to the depth of history taken into account. Given the limited recording lengths and for consistency with prior work [22], we set *k* = 1, which allows us to estimate the active information storage and the significance using the Pearson correlation coefficient (Eq. 4).

#### 2.7.2 Bivariate functional connectivity

The simplest model of the statistical dependencies between cells is the functional connectivity network, which represents links between cells with the correlation or mutual information between their calcium time series. For each pair of cells, we computed the mutual informations between the associated time series, and then subtracted off the expected mutual information computed from a set of 5000 circular-shifted nulls:

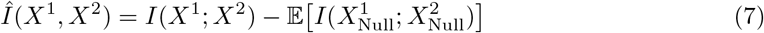

If *Î* (*X*^1^; *X*^2^) *<* 0 nat, it was set to zero, as there was no dependency not attributable to first-order autocorrelation. The resulting network represents only those cell-to-cell dependencies that exceed the baseline level of apparent connectivity that would be expected based solely on the properties of the individual time series.

#### 2.7.3 Higher-order interactions

To understand how groups of multiple cells interact, we used several multivariate generalizations of the mutual information (Eg. 3). These measures quantify different notions of multipartite interaction, and collectively provide a heuristic picture of how the emergent structure of the system changes in response to ATP.

For each of the records, we sampled 10,000 randomly selected triads (sets of three cells). From each triad we computed total correlation, dual total correlation, O-information, and S-information for each triad (all measures described below). We then corrected the apparent information by subtracting off the expected value of a null distribution constructed from 5,000 circularly-shifted nulls. We recorded two statistics: the density (i.e. what percentage of the 10,000 samples had corrected information ≠ 0), and the average corrected value for those triads that did show significant information.

##### Total correlation

The first measure is the total correlation [50] (independently derived by Tononi, Sporns, and Edelman as the “integration” [51]). The total correlation quantifies the extent to which a set of cells are coherently synchronized: it is zero in the case of global independence, and maximal in the case where each channel is a copy of every other one.

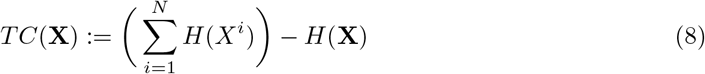

The total correlation can be understood as the Kullback-Leibler divergence of the true distribution from a prior of global independence:

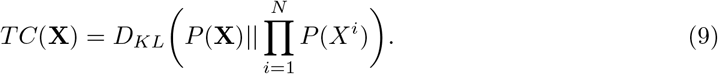

To correct for the possibility that first-order features of the data artificially inflate the total correlation, we correct the raw value with the expected value of a null distribution computed from circularly-shifted data:

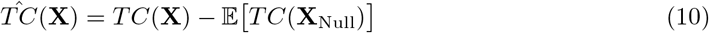

##### Dual total correlation

The second measure is the dual total correlation [52]. Unlike the total correlation which quantifies deviation from global independence, the dual total correlation quantifies how much information is shared over sets of two or more random variables:

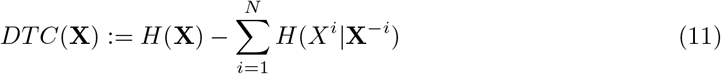

As with the total correlation, we correct the apparent dual total correlation with the expected value of the circular-shifted null:

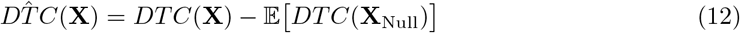

#### 2.7.4 Transfer entropy

The multivariate mutual informations assume that every frame is a draw from a multivariate Gaussian parameterized by covariance matrix Σ. While correcting for first-order autocorrelation partially addresses temporal dynamics, to more precisely characterize information “flows”, we used the transfer entropy [31, 27], which quantifies the degree to which knowing the *past* of one cell reduces our uncertainty about the next state of another cell after accounting for its own past. Formally:

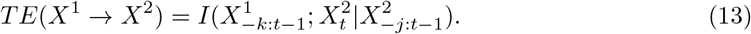

Here, the parameter *k* refers to the depth of history of *X*^1^ to account for and *j* refers to the depth of history of *X*^2^ to account for. We set *k* = *j* = 1 frame of history. While this is somewhat *ad-hoc*, the short recording times limit the dimensionality that is accessible without running into finite size effects.

As with before, we correct each transfer entropy by subtracting off the expected null transfer entropy from a distribution of 1,000 circularly-shifted nulls:

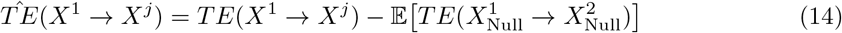

Once again, we record both the density (what proportion of edges have transfer entropy greater than 0), as well as the average transfer entropy across all non-zero edges.

#### 2.7.5 Integrated information

The bivariate transfer entropy only captures one aspect of time-directed information dynamics (information transfer). To understand how ATP alters dynamic information integration, we used the Φ^*W MS*^ measure [32]. For a system **X** partitioned into two subsystems: **X**_*α*_ *⊂* **X** and **X**_*β*_ = **X***/***X**_*α*_, the integrated information is the amount of predictive power about the future of the whole **X** that is not accessible when the “parts” (**X**_*α*_, **X**_*β*_):

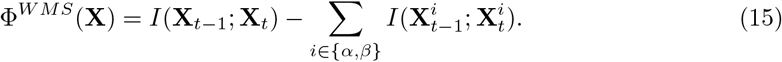

Intuitively, we can think of Φ(**X**) as quantifying the extent to which the “whole” is greater than the sum of its parts.

Unfortunately, the naive Φ measure can be negative. This occurs when there is substantial redundant information copied over both **X**_*α*_ and **X**_*β*_ [53]. To correct for this, Mediano et al., suggested a corrected measure that add back in that redundancy [54]:

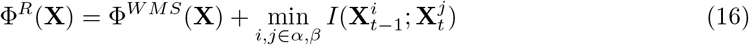

While the use of minimum mutual information as a measure of temporal redundancy has been critiqued [55], the Φ^*R*^ measure has been shown to be sensitive to different states in artificial and biological systems, reliably indexing level of consciousness in human brains [56], and relative complexity in artificial systems [33, 57].

Here, we computed Φ^*R*^ for every pair of cells in each bot and corrected the final value by subtracting off the expected value from a distribution of 1000 circular-shifted nulls.

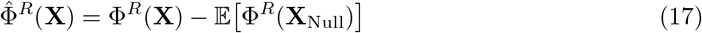

This approaches allows us to record both the density of pairs that integrate information and the average magnitude of the interaction for those pairs with non-zero Φ^*R*^.

### 2.8 Lempel-Ziv complexity

The final analysis was the Lempel-Ziv complexity of the calcium time series [58]. The Lempel-Ziv algorithm measures the compressibility of a dataset by constructing a dictionary of sub-strings that losslessly encode the original data in reduced form. Following prior work [59, 60, 61] we first binarized each signal about its mean, and then flattened the resulting binary array column-wise to produce a single, long binary string. This was input into the Lempel-Ziv algorithm, and the length of the resulting dictionary was recorded.

The size of the dictionary will be maximal when the original data is algorithmically random. To correct for the possibility of autocorrelation reducing the apparent randomness, each estimate was divided by the expected value of a null distribution composed of 5,000 circular-shifted nulls:

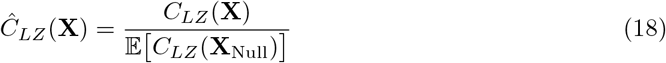

Unlike the measures described above, the LZC analysis does not rely on assumptions of Gaussianity - it is instead a non-parametric measure of the compressibility of the global patterns of calcium activity.

## 3 Results

### 3.1 ATP exposure reduces cellular long-term memory

To test how ATP altered the behavior of single-cells, we first computed the lag-1 active information storage (Eq. 6). This served two purposes: the first is to gain insight into the biological effects of ATP on individual cells. Secondarily, building estimators of higher-order, multicellular interactions requires understanding the lower-order behavior of the component “parts” and accounting for them (as in the null model construction). We found that exposure to ATP caused a significant decrease in the lag-1 autocorrelation for the individual calcium time series (Wilcoxon signed-rank: W=6, p=0.013, median percentage change = -13.92%). For visualization, see Figure 2A. This indicates that individual cells are responsive to ATP and that a major effect of the molecule is to increase the “noisiness” of calcium activity - effectively reducing the dependence of cellular activity on its own past. This change also justifies the use of the circular-shift null, as it is known that autocorrelation can artificially inflate measures of dependency [47]. By subtracting off autocorrelation-preserving nulls, we can correct for spurious change in dependency attributable to alterations to cell-level active information storage. For visualization, see Figure 2A.

**Figure 2:**
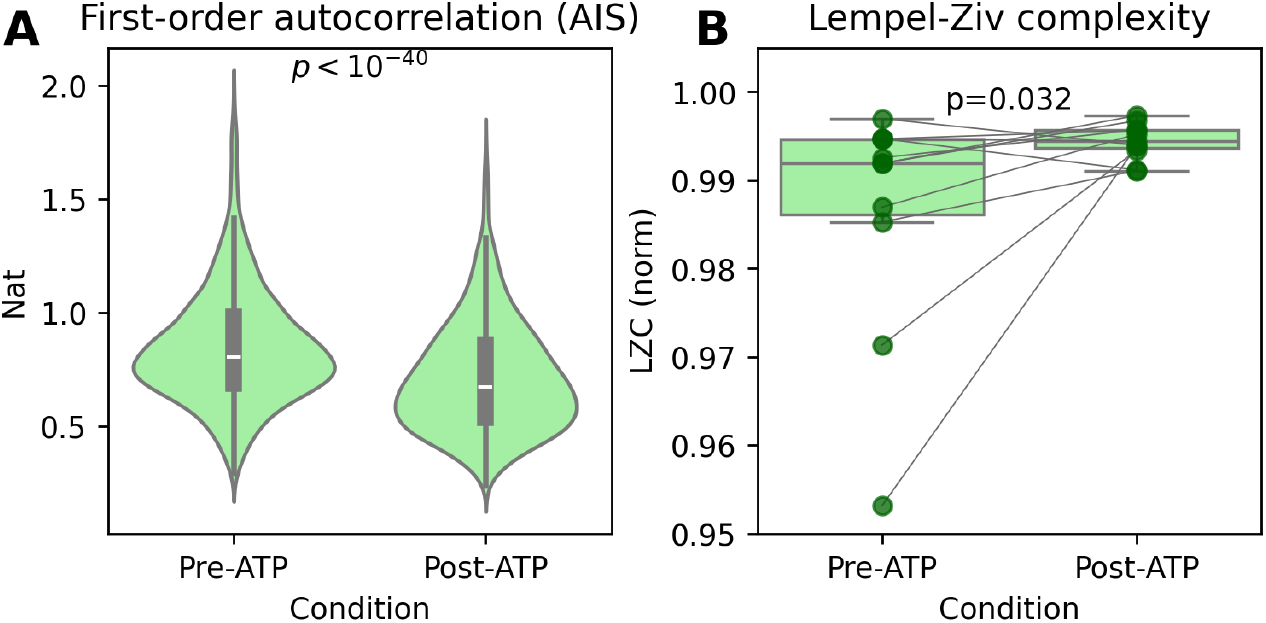
ATP alters entropy-rate dynamics at local and global levels. **A:** Exposure to ATP prompts a small but significant decrease in the lag-1 active information storage [28] for each cell individually (Wilcoxon signed-rank: W=6, p=0.013, median percentage change = -13.92%), indicating an increase in entropy rate. **B:** At the global level of all cells, the Lempel-Ziv complexity [59, 60] increases (Wilcoxon signed-rank test: W=9, p=0.03, median percentage change = 49.8%), indicating a macro-scale increase in entropy-rate.

### 3.2 ATP reduces pairwise and higher-order information structures

Having established that ATP alters the information-processing dynamics of individual cells, we then asked how exposure altered pairwise and higher-order coordination between sets of cells.

To do this, we first used the pairwise mutual information (Eq. 3) and then triadic total and dual total correlation (Eqs 8, 11). These measures provide an increasingly higher-order picture of multi-cellular organization across the surface of the basal Xenobot.

ATP exposure significantly reduced mean pairwise mutual information (Wilcoxon signed-rank test: W=5, p=0.0097, median percentage change = -29.58%, see Figure 3A). Curiously, while the global average pairwise mutual information decreased, the density (i.e. the percentage of possible edges with mutual information significantly greater than zero) did not change at all (Wilcoxon signed-rank test: W=17, p=0.174, see Figure 3B). Unfortunately, it is not possible to determine whether the preserved network density reflects stability of individual connections or balanced turnover, due to the absence of consistent cell identities across conditions.

**Figure 3:**
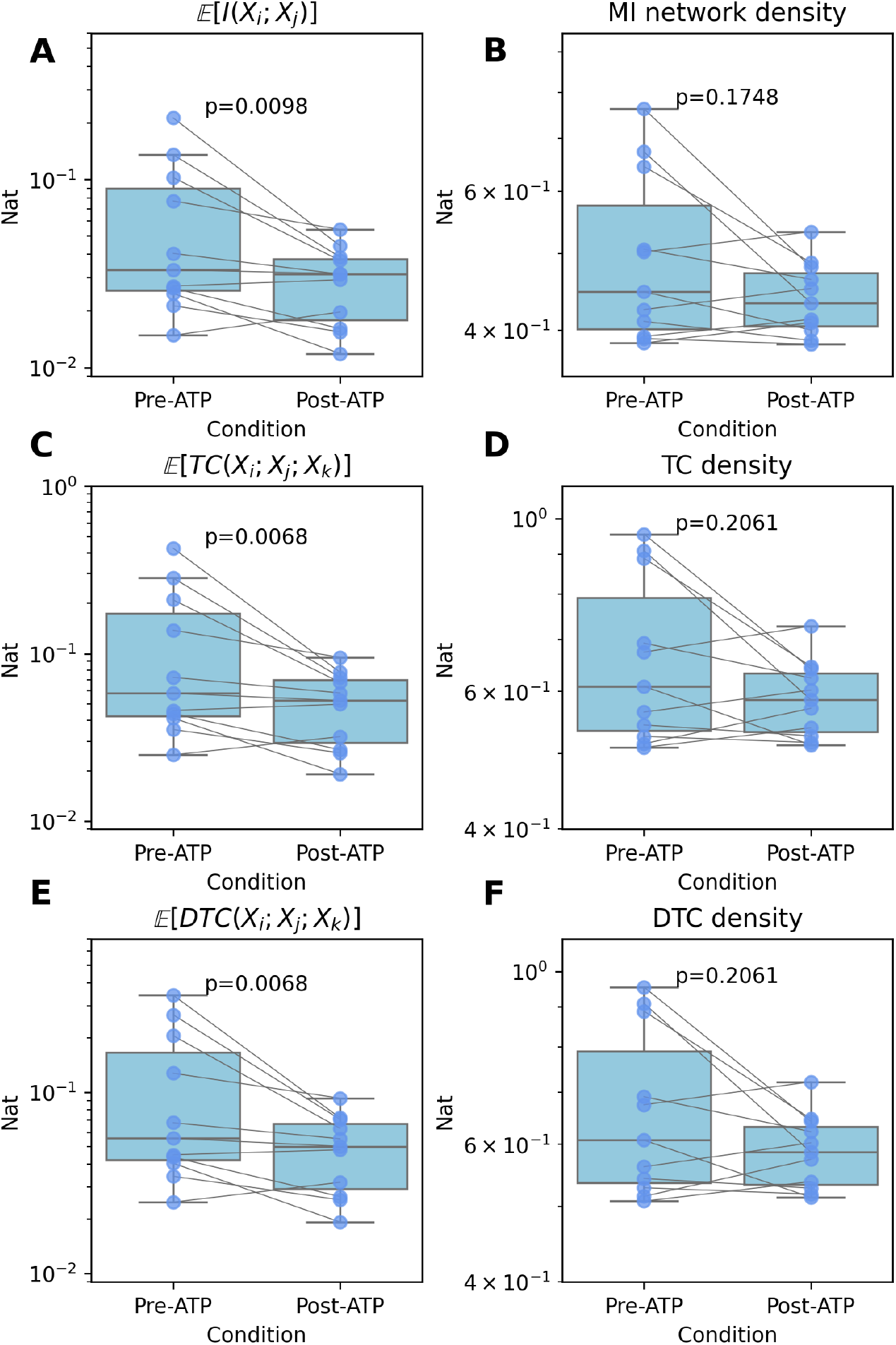
Pairwise and higher-order functional connectivity. **A:** 4 hours post exposure, pairwise functional connectivity had decreased, but **B:** the average network density (the proportion of possible edges with a mutual information greater than zero) had not changed. **C-F:** The same pattern was observed for triadic total correlation and triadic dual total correlation: for both measures, the information magnitude was significantly lower, the proportion of significant triads was not changed.

The next measure is the total correlation [50], which serves as a heuristic measure of higher-order redundancy [62] and quantifies the global deviation from independence. We found that ATP significantly decreased the average triadic total correlation (Wilcoxon signed-rank test: W=4.0, p=0.0068, median percentage change = -31.05%, see Figure 3C). There was no significant change in the proportion of sampled triads that showed significant total correlation, however (Wilcoxon signed-rank test: W=18, p=0.21). See Figure 3D. We also tested the dual total correlation, a heuristic measure of synergy that quantifies how much information is shared over sets of two or more elements. Like the total correlation, we found a similar decrease in information (Wilcoxon signed-rank test: W=4.0, p=0.0068, median percentage change = -28.71%, see Figure 3E). As with the pairwise functional connectivity, there was no difference in the triadic dual total correlation density (i.e. the proportion of possible triads with significantly non-zero dual total correlation): Wilcoxon signed-rank test: W=18, p=0.2, see Figure 3F. Collectively, these results suggest that the effect of ATP is not necessarily to simply silence functional channels (which would result in a decrease in network density), but rather, to decrease the degree of functional correlation between pairs of cells. In other words, hinder the communication and co-ordination across the cells of the basal Xenobots.

### 3.3 ATP reduces time-directed information flow and integration

Thus far, the primary measures of higher-order organization (mutual information, total correlation, and dual total correlation) have all be static measures: they model every frame as draw from some high-dimensional probability distribution, with no accounting for how past states inform on present ones. To explore temporal dynamics and gain a deeper understanding of how information flows through the Xenobots, we turned to the transfer entropy [30] (Eq. 13), which measures how the past of one cells informs on the future of another. ATP exposure reduced the average lag-1 transfer entropy [27] (Wilcoxon signed-rank test: W=3, p=0.0049, median percentage change = -13.61%, see Figure 4B), as well as a weak change in the density of the network (Wilcoxon signed-rank: W=10, p=0.042, median percentage change= -6.57%, see Figure 4C).

**Figure 4:**
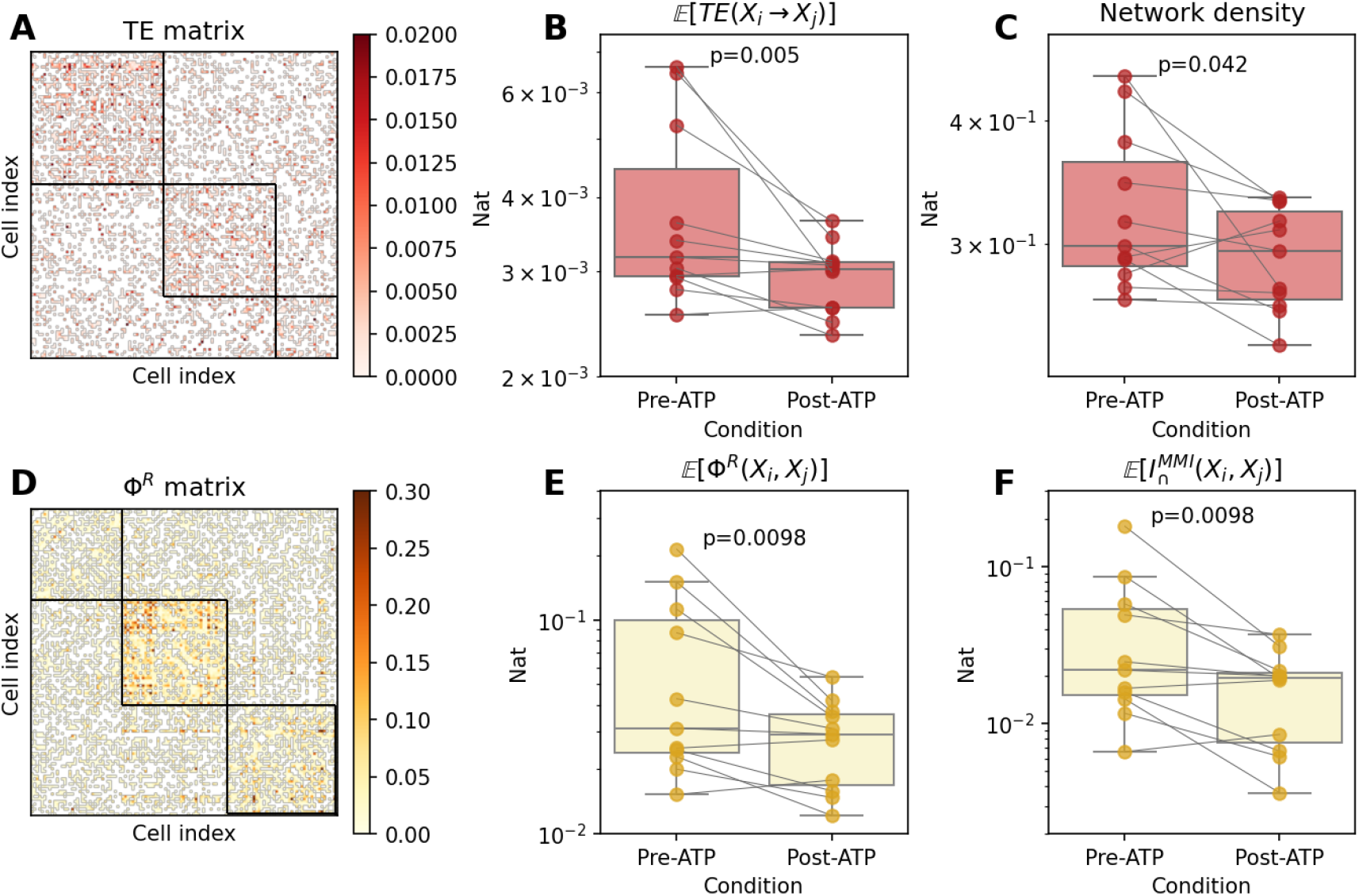
Temporal dynamics results. **A:** An example adjacency matrix showing edges with a significant lag-1 transfer entropy. Clustering was done with a modified multi-resolution consensus clustering algorithm purely for visual effect [49]. **B:** ATP exposure significantly decreased the average pairwise transfer entropy (Wilcoxon signed-rank test: W=3, p=0.0049, median percentage change = -13.61%) but **C:** but barely altered the density of the network overall (Wilcoxon signed-rank: W=10, p=0.042, median percentage change= -6.57%). **D:** the pairwise corrected integrated information matrix (Φ^R^), similarly clustered with the multi-resolution consensus clustering algorithm for visualization. This is the same basal Xenobot and same condition as in A. **E:** The average pairwise Φ^*R*^ for each culture before and after ATP. ATP exposure reduced global information integration significantly (Wilcoxon signed-rank test: W=5, p=0.0098, median percentage change = -35.78%). **F:** ATP exposure also reduced the global redundancy as well (given by the minimum mutual information) - Wilcoxon signed-rank test: W=5, p=0.0097, median percentage change = -46.89%.

The transfer entropy, like the pairwise mutual information is generally understood as a pairwise measure (although it does have a rarely-recognized higher-order component [63, 64]). To produce a genuinely higher-order measure of information flow, we used the corrected integrated information [33] (Eq. 16), which considers the irreducible information shared by four variables: two variables in the present and in the future. Like the transfer entropy, ATP also reduced the corrected integrated information Φ^*R*^ (Wilcoxon signed-rank test: W=5, p=0.0098, median percentage change = -35.78%, see Figure 4F), the density of the integrated information network (Wilcoxon signed-rank test: W=6, p=0.014, median percentage change = -10.34%), and the minimum mutual information redundancy (Wilcoxon signed-rank test: W=5, p=0.0097, median percentage change = -46.89%, see Figure 4F).

The final measure was the global Lempel-Ziv complexity [58]. Thus far, even the highest-order measures have been restricted to sets of three or four random variables, and therefore provided limited insight into the global structure of activity of the whole bot. The Lempel-Ziv complexity describes the compressibility of the whole multivariate time series, and in some sense provides the “highest”-order picture of spontaneous calcium dynamics before and after ATP exposure. ATP exposure increased the normalized Lempel-Ziv complexity (Wilcoxon signed-rank test: W=9, p=0.03, median percentage change = 49.8%, see Figure 2B). This result stands in contrast to the established pattern of changes reported above: functional and effective connectivity analyses generally reported a decrease in information flow between cells, but the Lempel-Ziv results show an increase in complexity/decrease in compressibility. These suggest that ATP is not simply “damping” the activity, but rather, globally “disorganizing” it, which is consistent both with the decreased cell-level autocorrelation, and the maintenance of network density.

## 4 Discussion

In this study, we found that exposing basal Xenobots to extracellular ATP caused dramatic reconfiguration of the functional and effective cellular information-processing networks within the bodies of the bots. Across a battery of measures, we found that eATP caused functional dependencies to get weaker (indexed by falling pairwise and higher-order functional connectivity measures) and more disorganized (indexed by changes in the Lempel-Ziv complexity of the calcium time series). Collectively, these results suggest that a primary effect of eATP exposure on basal Xenobots is to “fragment” them - individual cells are not necessarily less active following exposure, but they are more independent from their neighbors, and less predictable over time. This suggests that information flows across the system are not “hardwired” into groups of cells, but rather are fluid and amenable to changes in response to environmental/external factors. This flexibility may be useful in facilitating adaptability to different conditions, while also bringing with it vulnerability to being “hacked” or “programmed” by outside entities that can learn which stimuli lead to changes in information-processing architectures. This is particularly relevant in studies of cancers, which is hypothesized to occur when groups of cells have become “disintegrated” from the larger collective [65]. An empirical test of this hypothesis may be to use the techniques described here to probe how patterns of functional connectivity or information flow vary over the boundary between cancerous and non-cancerous neighboring cells.

One possible explanation for this phenomenon comes from studies of the “cell danger response” (CDR) [66]. The CDR is an ancient and well-conserved set of metabolic changes that occur in cells in response to the detection of a threat in the extracellular environment. The CDR is analogous to the classic fight/flight/freeze response to danger in animals, although since cells have limited capacity to fight or flee, it is most like the freeze state. Crucially sensing eATP in the environment is a reliable trigger for the CDR. When activated, the CDR sends cells into a defensive posture, altering metabolism (such as a shift from oxidative phosphorylation toward glycolysis) and reducing activity in mitochondria, which in turn increases intracellular oxidative load. It is possible that, in this defensive state, individual cellular survival comes at the expense of multicellular coordination - cells are, in a sense “defecting” from the collective to increase their own likelihood of surviving. Currently, our results cannot test whether the CDR itself is activated in these cells, but future work could test whether the degree of change in informational structure was correlated with expression of particular indicators of the CDR (such as altered purines, pyrimidines, sphingolipids, and phospholipids). Aberrant purinergic signaling has been implicated in the pathophysiology of a number of chronic diseases, including myalgic encephalomyelitis (also known as chronic fatigue syndrome) [67, 68], Gulf War illness [69], as well as developmental disorders such as autism spectrum disorders [70, 71]. Consequently, the results presented here may provide insight into a common underlying feature of all these illnesses: namely a breakdown of the multi-cellular coordination required to support higher-order emergent structures.

While basal Xenobots are strictly non-neural organoids, and eATP is usually considered a neuromodulator, it is interesting to compare the effects observed here with psychopharmacological interventions in the brain, as such comparisons may help us better build an intuitive understanding of what these changes might mean. The pattern observed here (reduced overall information integration coupled with an increased Lempel-Ziv complexity) is somewhat unusual. Generally, reductions in the information-density of neural activity are associated with reductions in level of consciousness [72, 73, 24, 74, 75], and this typically includes Lempel-Ziv complexity as well [76, 59, 61]. In contrast, an increase in Lempel-Ziv complexity is typically associated with “consciousness expanding” drugs such as psychedelics [60, 77, 78], or meditative states [79, 80], where the effects on global information flow are variable. This suggests that the effects of ATP on information-processing dynamics in basal Xenobots do not cleanly map onto the usual landscape of psychopharmacological interventions and may instead represent a novel kind of of (dys)regulation. Future research exploring other compounds (such as antipurinergics like suramin, or other signaling compounds) may reveal a novel space of interactions between biological signals and self-organizing information structures.

Finally, these results have implications for bioengineering, regardless of what the underlying mechanism might be. It appears that one may be able to use purinergic signaling to “tune” the overall degree of “emergent integration” in biological tissue, moving up and down a gradient from “independent cells” to “integrated organismal whole.” More generally, these results represent a proof-of-concept that “emergent engineering” in living systems is a viable research program. The current demonstration, while limited in scope represents a novel avenue by which micro-scale interventions may be leveraged to produce macro-scale change self-organization and information-integration capacity. For engineers building with active matter, the ability to control the overall degree, or pattern of, information integration within a system represents a powerful design parameter. Recent work in artificial intelligence systems has shown that modulating over-all information integration can improve the performance on a variety of tasks [81, 82, 83, 84], and in neuroscience, emergent informational scaffolds have been shown to associate with cognitive performance over the course of infant development [85]. In the context of bioengineering, similar capabilities may be discovered in the context of regeneration, healing, and cancer therapeutics.

## 5 Conclusions

We have shown that exposure to extracellular ATP disrupts the emergence of coordinated, multi-cell information-processing networks. A battery of functional and effective statistics shows a consistent pattern of decreased multi-cellular coordination, more unpredictable temporal dynamics, and a global disorganization of activity. These results suggest that purinergic signaling could serve as a control knob for bioengineers and scientists to explore the physical substrates that underpin the emergence of macro-scale phenomena in biology. More generally, this work lays the foundation for a general capacity to change the structure of information integration systems in non-neural, multi-cellular collectives. The current focus on ATP is a proof-of-principle that can be further explored and opens up a research program at the intersection of multiple fields, including evolutionary biology, bioengineering, complex systems, and cognitive science.

## Acknowledgments

M.L. gratefully acknowledges support of the John Templeton Foundation (via grant 62212). The opinions expressed in this publication are those of the authors and do not necessarily reflect the views of the John Templeton Foundation. TFV is supported by the Cold Regions Research and Engineering Laboratory (CRREL) under Contract No. W913E524C0012

## Author contribution statement

V.P. performed the eATP experiments and collected the calcium imaging data. T.F.V. processed the videos, performed the analyses, and wrote the initial draft. V.P., M.L. and J.B. provided edits for the manuscript.

## Data availability statement

CSV files of the calcium series will be available as supplemental information upon final publication.

